# Integration of iPSC-derived microglia into human midbrain organoids enhances microglial maturation and inflammatory signaling

**DOI:** 10.64898/2026.04.06.716748

**Authors:** Emma J. MacDougall, Ghislaine Deyab, Apoline Ormancey, Jialun Li, Taylor M. Goldsmith, Paula Lépine, María Baeza Trallero, Nadja Finkel, Julien Sirois, Martin H. Berryer, Thomas M. Durcan, Edward A. Fon

**Affiliations:** Neurodegenerative Disorders Research Group, Department of Neurology and Neurosurgery, Montreal Neurological Institute-Hospital (The Neuro), McGill University, Montreal, H3A 2B4, Canada; Early Drug Discovery Unit, Montreal Neurological Institute-Hospital (The Neuro), McGill University, Montreal, H3A 2B4, Canada

## Abstract

Microglia are the resident immune cells of the central nervous system and play key roles in the healthy brain during development and adulthood, as well as during neurodegenerative diseases - including Parkinson’s disease (PD). Yet the role of microglia in PD pathogenesis has not been fully elucidated. Limitations of 2D cell culture and animal models in simulating human microglia in the brain parenchyma have contributed to this knowledge gap. Human midbrain organoids (hMOs) provide a promising model that can recapitulate elements of PD pathology but lack microglial cells. Here we adapt protocols for the differentiation of hMOs and human iPSC-derived microglia (iMG) to generate iMG-hMO assembloids. Within assembloids, integrated iMG (intMG) express canonical microglia markers and induce the release of cytokines and chemokines. Transcriptomic profiling by single cell RNA sequencing reveals that intMG adopt a more mature and inflammation-responsive state compared to 2D iMG. The integration of microglia results in increased signaling through inflammatory and trophic pathways that drive altered transcriptional signatures of dopaminergic neurons and astrocytes within assembloids. Overall, iMG-hMO assembloids have the potential to more faithfully model the role of microglia and neuroinflammation in PD pathogenesis.

## Introduction

Microglia are the tissue-resident immune cells of the central nervous system (CNS). They arise from early erythromyeloid precursors that migrate from the yolk sac to the developing CNS,^1-3^ and in adulthood account for approximately 0.5-16% of brain cells.^4^ Microglia are highly dynamic and can adopt diverse, multidimensional functional states - marked by morphologic and transcriptomic, among other changes - in response to their environment. Surveillance, phagocytosis, and release of soluble factors are core microglial functionalities; allowing microglia to play roles in neurogenesis and synapse remodeling, trophic support, tissue repair, inflammation, and clearance of exogenous pathogens and endogenous cellular debris.^5^ Microglia are increasingly regarded as important in the etiology of and response to neurodegenerative diseases, including PD.^6-9^ However, it remains unclear if the changes in microglial state and functionality observed in PD are beneficial or detrimental, triggered by or contribute to neuronal death, and if this paradigm shifts during disease course. A better understanding of the contribution of microglia to PD pathogenesis is necessary for guiding development of proposed immune-targeting therapies for PD.^10^

Although microglia share a conserved set of core genes and proteins across species, transcriptomic and proteomic divergence of human microglia is becoming more apparent.^11-14^ Historically, studying human microglia has been very challenging; access to human primary microglia is rare and culturing these cells *in vitro* has been shown to lead to extensive transcriptional changes.^15,16^ Human microglia-like cells derived from induced pluripotent stem cells (iPSCs) are a renewable resource that can be derived from patients with neurological diseases and are amenable to genetic engineering.^17-22^ iPSC-derived microglia (iMG) have been co-cultured with 2D iPSC-derived neurons,^23,24^ integrated into brain organoids to form assembloids,^25^ and xenotransplanted into mouse brains to model the interactions of microglia with other neural cells, and mitigate the effect of *in vitro* culture on microglial transcriptional state.^17,26-28^ Of these, only brain organoids provide a fully human, complex 3D microenvironment to model brain function and brain disease. However, as brain organoids generally lack endogenous myeloid-lineage cells, namely microglia,^25^ different approaches have been taken to introduce microglia into brain organoids.^17,29-36^ These assembloid models have been shown to induce both *in vivo*-like transcriptional signatures in microglia,^31,36^ and increased neuronal and network maturation.^29-36^

Most studies using such assembloids employed either cerebral or cortical organoids, which lack the regional specificity to model PD. In contrast, human midbrain organoids (hMOs) model the brain region most affected by dopamine neuron degeneration in PD and have been shown to recapitulate key features of PD pathology.^37-40^ Here, we adapted and optimized a recently published protocol,^41^ to characterize iMG-hMO assembloids (from here on referred to as assembloids) and their potential to model PD. We confirm microglial integration, maturation, and functionality within assembloids. Further, single-cell RNA sequencing reveals that iMG that are integrated (intMG) within assembloids have a more mature and inflammation-responsive transcriptional signature compared to their 2D counterparts. Finally, although integration of microglia does not lead to alterations in the number of DA neurons or astrocytes in assembloids, it led to changes in their transcriptional signatures and predicted signaling patterns.

## Results

### iPSC-derived hematopoietic precursor cells integrate and mature into microglia within assembloids

Assembloids were generated by adapting previously published protocols for the generation of hMOs, the production of iPSC-derived hematopoietic precursor cells (iHPCs) and the differentiation of iHPCs to 2D iMGs.^18,42,43^ hMOs were generated from iPSCs derived from a healthy control donor and cultured for two months. iHPCs differentiated from the same control iPSC line were then added into the same vessel at a ratio of 50 000 iHPCs per hMO. At this time, the culture media was changed from hMO maturation media to assembloid media consisting of microglial differentiation media with the addition of hMO growth and maturation factors brain derived neurotrophic factor (BDNF), glial derived neurotrophic factor (GDNF), and ascorbic acid (AA). Dibutyryl-cyclic AMP (db-cAMP), a component of hMO media, was excluded due to its toxicity to 2D iMGs as demonstrated by Sabate-Soler and colleagues.^41^ iHPCs and hMOs were cultured together for an additional month in assembloid media to allow integration and maturation of iHPCs into intMG (Fig. 1a). For comparison, iHPCs from the same batches used to generate assembloids were differentiated into iMGs in 2D, and hMOs were maintained in assembloid media without the addition of iHPCs. Immunofluorescence (IF) staining confirmed integration of iHPCs into assembloids, and differentiation into intMG as substantiated by the expression of the canonical microglial marker Iba1. The total area of Iba1 staining and number of Iba1 positive cells was significantly increased in assembloids, compared to hMOs (Fig. 1b-d). Tissue clearing followed by 3D imaging of assembloids stained for Iba1 further illustrates the infiltration of intMG and the adoption of classical ramified microglial morphology (Fig. 1e, Supplementary movies 1-5). The ongoing maturation of iHPCs into intMG was further supported by expression of microglial transcription factor PU.1 and macrophage marker CD68 in Iba1 positive cells in assembloids but not in hMOs, as detected by IF staining (Fig. 1f-g). To assess intMG survival, assembloids were maintained for an additional three months after addition of iHPCs. The presence of intMG at this later time-point was confirmed by Iba1 IF staining in assembloids but not hMOs cultured in parallel (Fig. 1h). Ultimately, we observe integration of iHPCs into hMOs to form assembloids, and ongoing maturation of these iHPCs into intMG based on expression of canonical microglial markers.

**Figure 1.**
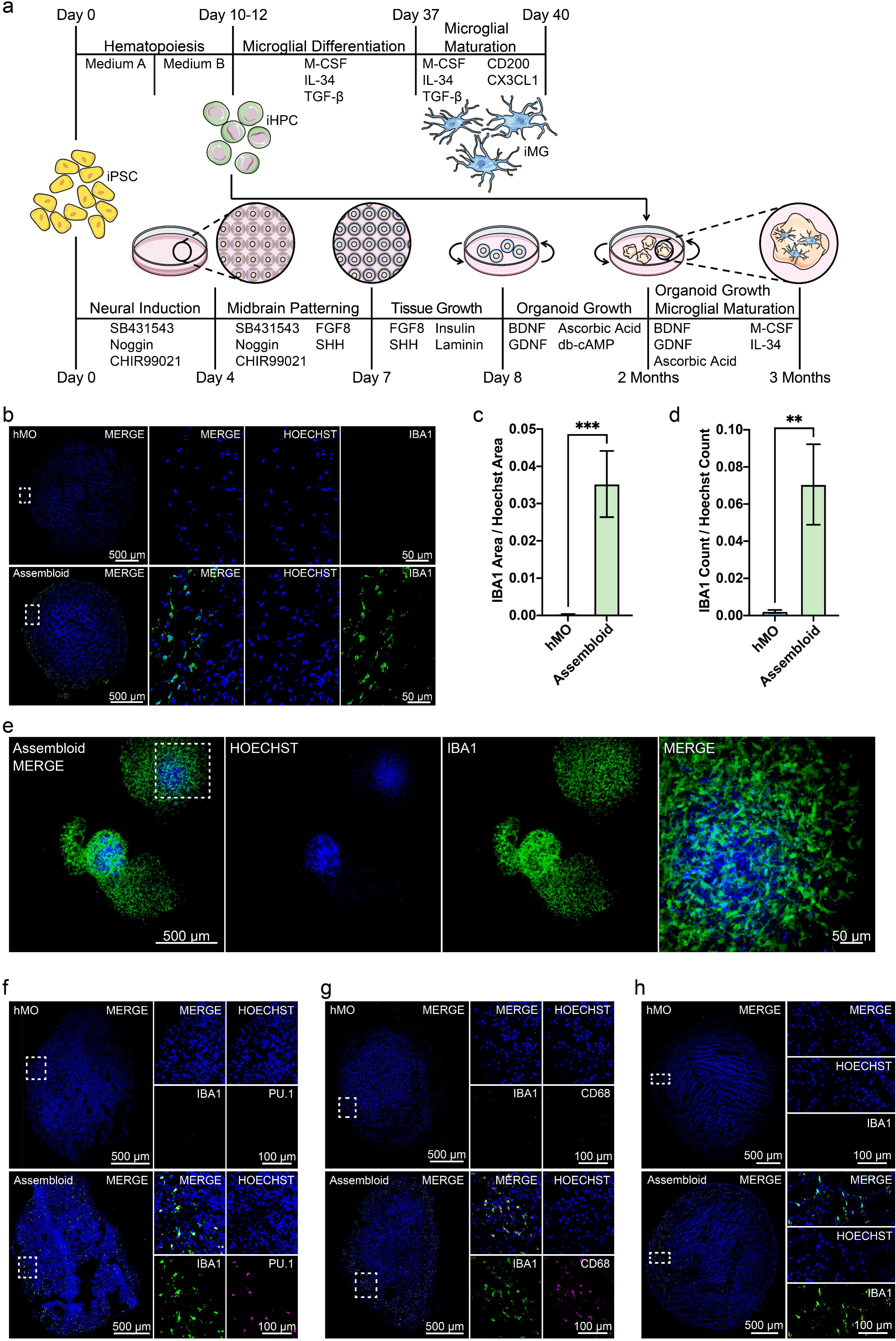
intMG colonize and mature within assembloids. **a** Schematic of iMG, hMO and assembloid generation. **b** IF staining of IBA1 in hMO and assembloids. **c** quantification of IBA1 positive area over Hoechst area, and **d** quantification of IBA1 positive cell count over Hoechst count in IF staining. **c-d** n = 3 batches, total 12 hMOs, 9 assembloids. Unpaired t-test * p ≤ 0.05, ** p ≤ 0.01, *** p ≤ 0.001, **** p ≤ 0.0001 **e** 3D IF imaging reveals ramified microglial morphology in assembloids. **f** IF staining of IBA1 and PU.1 and **g** IBA1 and CD68 in hMO and assembloids. **h** IBA1 expressing intMGs survive up to 5 months in assembloids, as shown by IF staining.

### Integrated microglia drive assembloid cytokine expression and secretion

It is well established that microglia increase the expression of cytokines and other soluble factors in response to proinflammatory stimuli. Thus, we treated assembloids, hMOs, and 2D iMGs for 24 hours with 100 ng/mL of lipopolysaccharide (LPS), a major component of the outer membrane of gram-negative bacteria and toll-like receptor 4 (TLR4) agonist known to trigger inflammatory signaling. mRNA expression levels of classical pro-inflammatory cytokines tumor necrosis factor α (*TNFα*), interleukin 1β (*IL-1β*), and interleukin 6 (*IL-6*) were then assessed via reverse transcription quantitative PCR (RT-qPCR) and represented as expression fold change in LPS versus non-treated samples (Fig. 2a). Expression of *TNFα* and *IL1β* were significantly increased, and there was a trend towards increased *IL-6* expression, in 2D iMGs and assembloids but not in hMOs in response to LPS treatment, indicating that intMG drive increased cytokine expression in assembloids. Induction of reactive astrocyte states in response to microglial signaling is marked by morphological and transcriptomic changes, including increased expression of glial fibrillary acidic protein (*GFAP*).^44-46^ We observe a significant increase in GFAP mRNA expression in response to LPS treatment in assembloids but not in 2D iMGs or hMOs; reflecting astrocyte reactivity, driven by intMG in response to LPS stimulus.

**Figure 2.**
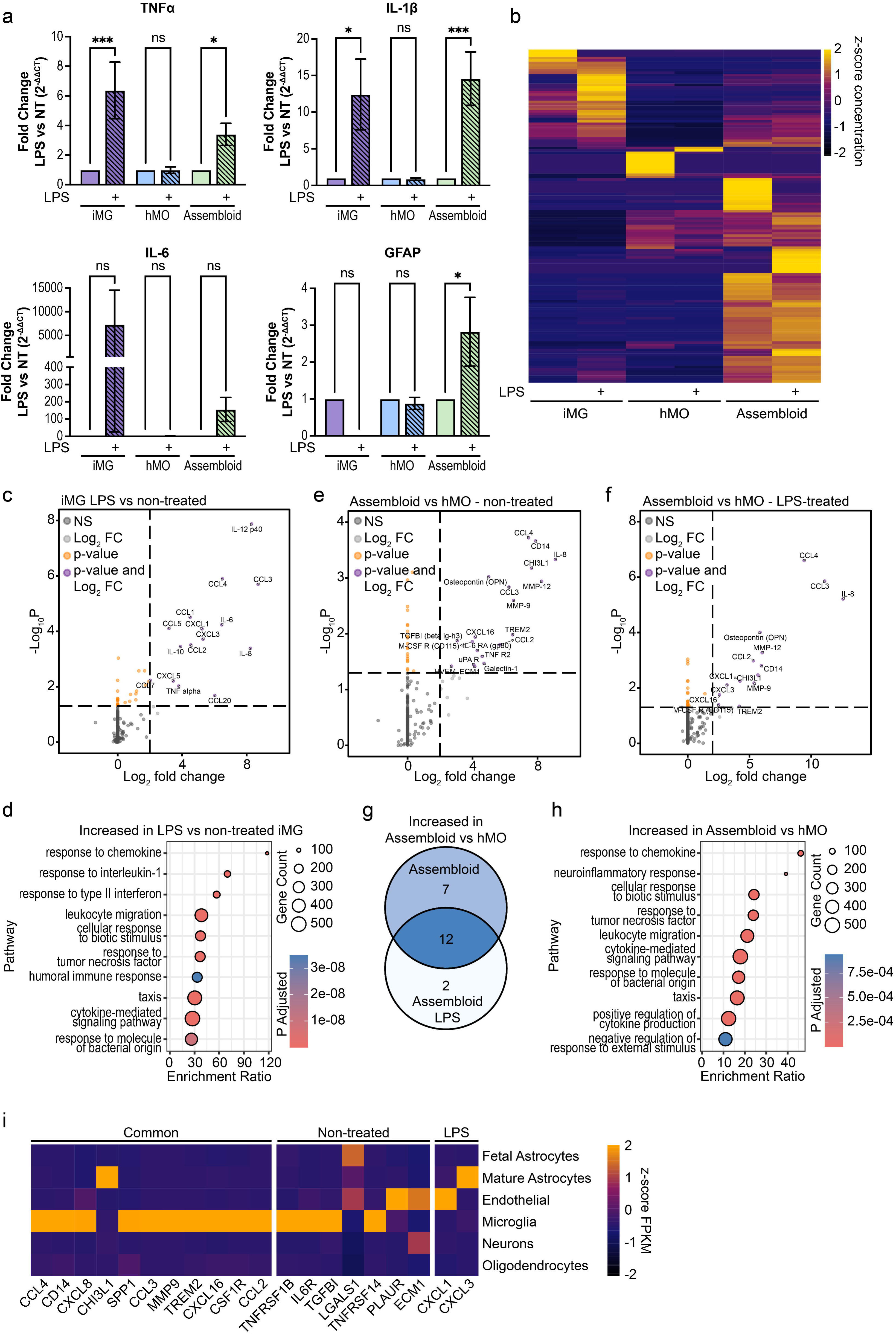
intMG drive cytokine and chemokine expression within assembloids. **a** RT-qPCR analysis of TNFα, IL1β, IL-6, and GFAP expression in 2D iMG, hMO, and assembloids nontreated or treated with 100 ng/mL LPS for 24 hrs. Expression is normalized to non-treated controls. n = 2 batches iMGs. n = 4 batches hMO/assembloids. Two way ANOVA with Šídák’s multiple comparisons test * p ≤ 0.05, ** p ≤ 0.01, *** p ≤ 0.001, **** p ≤ 0.0001 **b** z-score of concentration (pg/mL) of proteins detected in conditioned media from 2D iMG, hMO, and assembloids nontreated or treated with 100 ng/mL LPS for 24 hrs, as detected by nELISA. n = 2 batches iMGs. n = 4 batches hMO/assembloids. **c** Significant DSPs in LPS-treated 2D iMGs versus non-treated 2D iMGs. **d** GO terms associated with DSPs increased in 2D iMGs after LPS treatment. **e** Significant DSPs in non-treated assembloid versus non-treated hMO. **f** Significant DSPs in LPS-treated assembloid versus LPS-treated hMO. **c, e-f** Log2FC cut-off = 2, p-value cut-off = 0.05 student’s t-test. **g** Venn diagram illustrating common and treatment-specific DSPs in assembloids. **h** GO terms associated with common DSPs increased in assembloids regardless of treatment. **i** z-score of expression (FPKM) of assembloid DSPs in brain RNA sequencing data base.

To investigate the release of soluble factors from assembloids more broadly and more directly at the protein level, conditioned media from non-treated and LPS treated 2D iMGs, hMOs, and assembloids was subjected to multiplexed nELISA.^47^ Of 200 proteins assayed, a total of 137 different proteins were detected across all samples. Comparison of the mean concentrations of these proteins in conditioned media illustrates unique patterns of protein secretion across 2D iMGs, hMOs, and assembloids (Fig. 2b). We then performed pairwise comparisons to determine which proteins were significantly differentially secreted (DSPs). In 2D iMGs, secretion of 15 cytokines and chemokines, including classic pro-inflammatory cytokines TNFα and *IL-6*, was significantly increased under LPS stimulation (Fig. 2c). Gene ontology (GO) over-representation analysis of these DSPs returns terms related to immune response and cell taxis (Fig. 2d). Within hMOs and assembloids there were no significant DSPs in response to LPS treatment. However, between hMOs and assembloids there were 19 and 14 proteins whose secretion was increased in assembloids, under non-treated and LPS treated conditions, respectively (Fig. 2e-f). The majority of these DSPs are cytokines and chemokines. However, release of additional proteins, some related to extracellular matrix (ECM) maintenance (ECM1, matrix metalloprotease (MMP)-9, MMP-12), is also increased in assembloids compared to hMOs. Secretion of 12 proteins was increased in assembloids compared to hMOs regardless of LPS treatment (Fig. 2g). Secretion of CCL3, CCL4, and IL-8, all shown to be LPS-responsive in 2D iMGs, was increased approximately six to eight-fold in non-treated assembloids and approximately ten-fold in LPS treated assembloids. These results indicate that the identity of DSPs in assembloids is shared across non-treated and LPS treated conditions, but the magnitude of increase in secretion is larger, although not significantly, under LPS treatment. GO analysis of the 12 DSPs significantly increased in assembloids regardless of treatment illustrates the increased secretion of components of immune, neuroinflammatory, and cell taxis pathways, upon microglial integration (Fig. 2h). To predict whether increased secretion was driven by protein expression and release from intMG, or by induction of expression and secretion in other cell types following microglial integration, we queried a human brain RNA sequencing dataset (brainrnaseq.org).^48^ Microglia displayed the highest mean expression for the majority of DSPs (Fig. 2i). Of note, a number of DSPs (CHI3L1, LGALS1, PLAUR, ECM1, CXCL1, and CXCL3) have highest expression in astrocytes or endothelial cells. Together, our findings indicate that the majority of changes in protein secretion in assembloids are driven by expression in and secretion from intMG, recapitulating a key microglial functionality. Further, secretion of these soluble factors likely leads to downstream changes in additional cell types in assembloids, illustrated by increased GFAP expression and secretion of proteins typically expressed by astrocytes and endothelial cells.

### Integrated microglia display a more mature and inflammation-responsive transcriptional signature compared to 2D microglia

To more deeply assess the impact of differentiation of iHPCs into intMG within the 3D assembloid environment, as well as the impact of intMG on additional assembloid cell types (neurons and macroglia), we performed single cell RNA sequencing (scRNAseq) of 2D iMGs, hMOs, and assembloids. Low quality cells and putative doublets were excluded, Louvain clustering was performed, and uniform manifold approximation and projection for dimension reduction (UMAP) was applied to visualize cell clustering. Initially, three distinct cell clusters were identified. Based on expression of microglial markers and sample identity we annotated these clusters as 2D microglia, intMG, and neurons and macroglia (Fig. 3a-c). The 2D microglia cluster is found only in the iMG sample (Fig. 3b), and expresses microglial marker genes *AIF1* (IBA1), *SPI1* (PU.1), *P2RY12*, *CX3CR1*, *C1QA*, *TYROBP*, and *TREM2* (Fig. 3c). The neurons and macroglia cluster is found in both the hMO and assembloid samples (Fig. 3b) and does not express any microglial marker genes but does express neuronal marker gene *STMN2* (Fig. 3c). Finally, the intMG cluster is found only in the assembloid sample (Fig. 3b) and expresses microglial marker genes (Fig. 3c).

**Figure 3.**
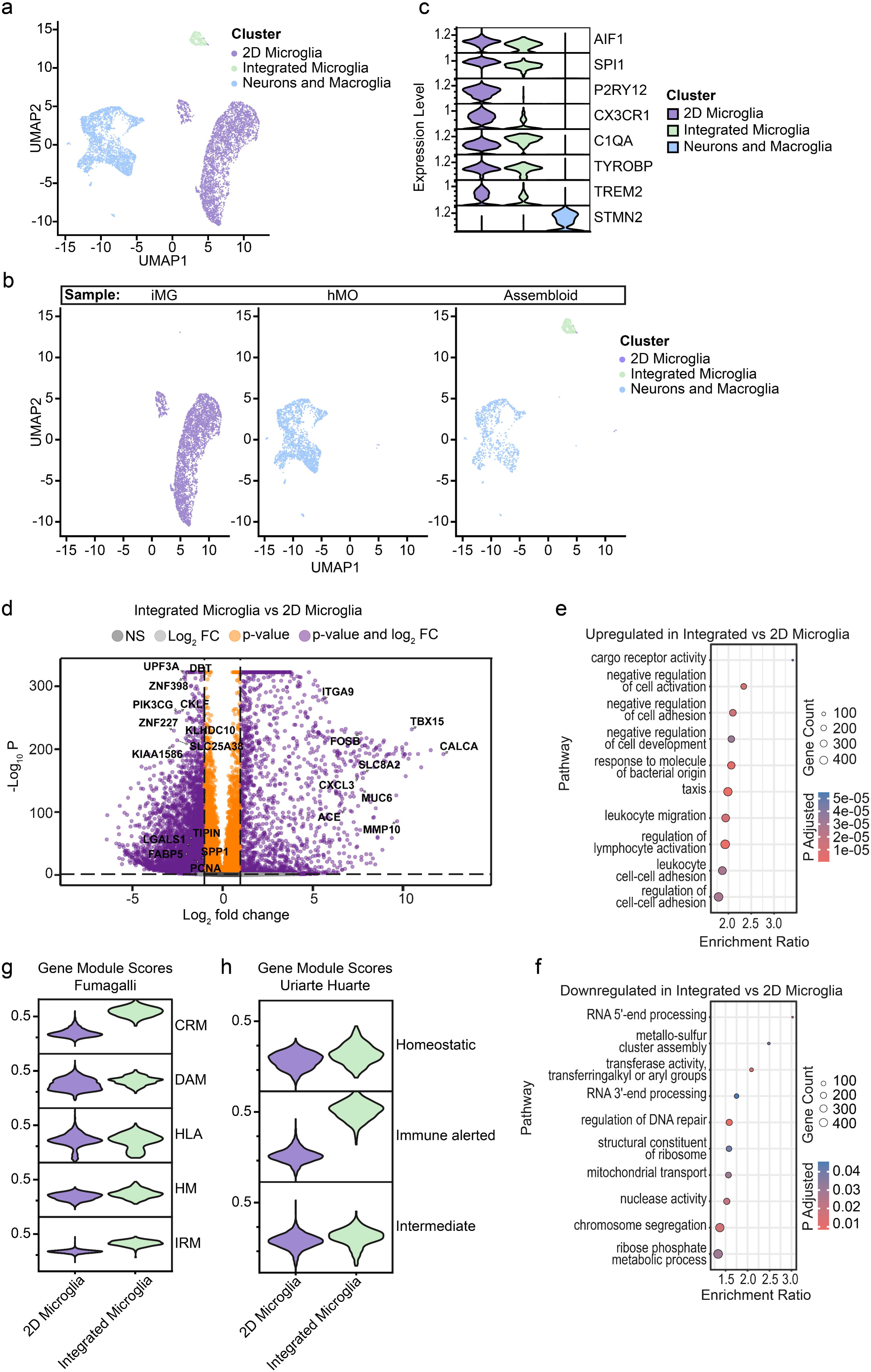
intMG display a distinct transcriptional identity compared to 2D microglia. **a** UMAP representation of scRNAseq data from 2D iMG, hMO, and assembloid cells shows three cell clusters. **b** Expression of microglial markers (AIF1, SPI1, P2RY12, TMEM119, CX3CR1, C1QA, TYROBP, TREM2) and neuronal marker STMN2 across the 2D microglia, intMG, and neuron and macroglia clusters. **c** UMAP representation illustrating the clusters present across samples. **d** DEGs up and downregulated in the intMG versus 2D microglia clusters. Log2FC cut-off = 1, p-value cut-off = 0.05 **e** GO terms associated with DEGs increased in intMG versus 2D microglia. **f** GO terms associated with DEGs decreased in intMG versus 2D microglia. **g** Gene module scores illustrate degree of similarity between 2D microglia and intMG gene expression and HM, DAM, CRM, IRM, and HLA reference gene sets defined by Fumagalli and colleagues. **h** Gene module scores illustrate degree of similarity between 2D microglia and intMG gene expression and homeostatic, immune-alerted, and intermediate reference gene sets defined by Uriarte Huarte and colleagues.

Pairwise comparison of microglia differentiated in the 3D assembloid environment revealed 2,153 genes that were upregulated, and 4,119 genes that were downregulated compared to microglia differentiated in 2D (Fig. 3d). scRNAseq profiling of microglia across the mouse lifespan has shown highly proliferative microglial states in embryonic and early post-natal brain. These early proliferative microglia are marked by expression of a number of genes involved in cell metabolism, proliferation, growth, and motility; including *FABP5*, *SPP1*, *LGALS1*, *TIPIN*, and *PCNA*^49^ – all of which are significantly downregulated in the intMG compared to 2D microglia cluster (Fig. 3d). GO analysis of the differentially expressed genes (DEGs) upregulated in the intMG compared to 2D microglia cluster returned terms related to regulation of cell activation and adhesion (Fig. 3e). Further, GO analysis of the DEGs downregulated in the intMG compared to 2D microglia cluster illustrates a decrease in expression of genes involved in regulating RNA and ribosome processing, cell cycle, and proliferation (Fig. 3f). Fumagalli and colleagues have recently identified five major transcriptional programs that have been reproducibly identified across multiple human microglial datasets. These transcriptional states include homeostatic microglia (HM), disease-associated microglia (DAM), antigen presenting microglia (HLA), interferon-responsive microglia (IRM), and cytokine-responsive microglia (CRM).^50^ Calculation of gene module scores illustrates an enrichment of the CRM and IRM signatures in intMG (Fig. 3g). Further, intMG display increased similarity to an immune alerted state found by Uriarte Huarte and colleagues to be enriched in the mouse midbrain (Fig. 3h).^51^ Taken together, our data indicates that differentiation of microglia within the 3D assembloid environment imparts a distinct transcriptional signature compared to those differentiated in 2D. intMG appear to be more mature than 2D microglia, evidenced by downregulation of genes in involved in proliferation and known to be expressed in embryonic and early post-natal mouse microglia.^49^ Further, intMG have increased expression of genes associated with inflammatory transcriptional programs, and pathways involved in cell activation and motility; likely reflective of inputs (signaling molecules, DAMPs, physical interaction, etc.) from other cell types in the assembloid.

### Assembloids do not contain altered proportions of neuronal or macroglial cells

We next assessed the impact of microglial integration on the neurons and macroglia within assembloids compared to those within hMOs. We extracted a subset of our scRNAseq data corresponding to cells arising from hMO and assembloid samples, integrated, re-clustered, and imputed this subset, resulting in 15 cell clusters. The clusters were annotated based on expression of characteristic cell type markers, described in detail in the methods section (Fig. 4a-b). We noted a subset of cells adjacent to the microglia cluster arising from hMO samples that did not express any microglial marker genes, but rather expressed fibroblast markers. To address this, these cells and the microglia cluster were isolated, subclustered into five clusters, and expression of microglial and fibroblast marker genes was assessed (Fig. S1a-c). Cluster 4 was exclusive to hMOs and expressed fibroblast markers *COL15A1*, *COL3A1*, and *MGP*. Thus cluster 4 was annotated as fibroblast cluster 2, while clusters 0-3 were annotated as microglia. In the brain, fibroblasts are a border enriched cell population that arise in part from both the mesoderm and neuroectoderm, and play an immunomodulatory role in response to injury.^52,53^ In hMOs, the absence of microglia may lead fibroblasts to adopt a more immunomodulatory state, underlying the close clustering of the fibroblast 2 cluster to microglia in our scRNAseq data. Apart from the Fibroblast 2 cluster and microglia cluster, which were unique to hMO and assembloid samples respectively, there were no significant differences in the proportions of the additional clusters in assembloids compared to hMOs as assessed by permutation testing (Fig. 4c-d). Together, these findings indicate that the integration of microglia into assembloids does not lead to any marked changes in the proportions or identities of the majority of neuronal and macroglial cell types.

**Figure 4.**
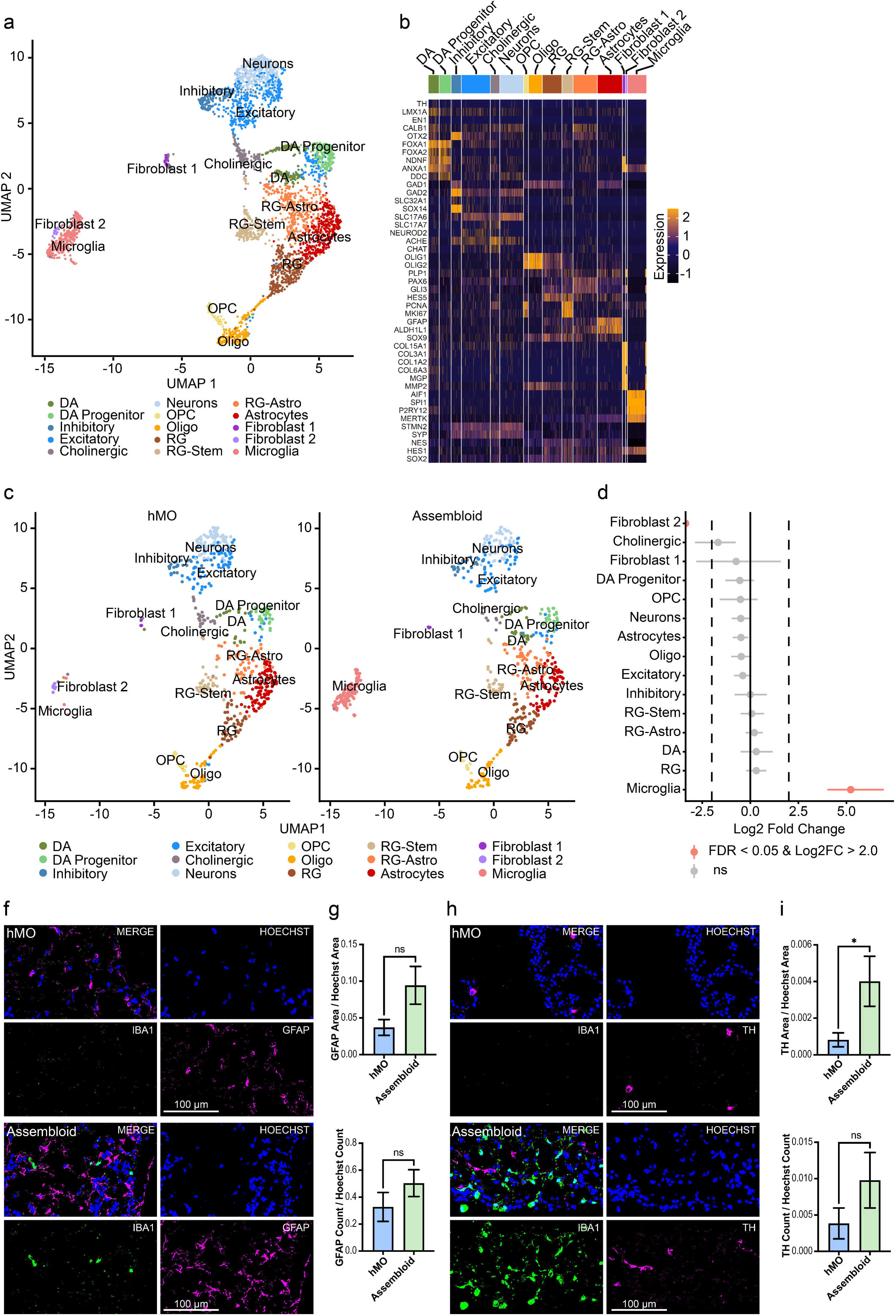
Assembloids do not have altered proportions of astrocytes and DA neurons. **a** UMAP representation of scRNAseq data from hMO and assembloid shows 15 cell clusters. **b** Expression of classical marker genes delineates cell clusters. **c** UMAP representation illustrating clusters present in hMOs and assembloids. **d** Quantification of cell clusters in assembloids versus hMOs. Log2FC cut-off = 2, p-value cut-off = 0.05. p-values calculated by permutation test. **f** IF staining of GFAP in hMOs and assembloids. **g** Quantification of GFAP positive area over Hoechst area and quantification of GFAP positive cell count over Hoechst count in IF staining. **h** IF staining of TH in hMOs and assembloids. n = 3 batches, total 8 hMOs, 9 assembloids. **i** Quantification of TH positive area over Hoechst area and quantification of TH positive cell count over Hoechst count in IF staining. n = 3 batches, total 12 hMOs, 9 assembloids. **g, i** Unpaired t-test * p ≤ 0.05, ** p ≤ 0.01, *** p ≤ 0.001, **** p ≤ 0.0001

Microglia – brain organoid assembloids have been shown to have increased neuronal maturation, reflected by changes in electrical activity, and altered expression of genes involved in synapse and DA circuit formation.^30-34,41^ Moreover, microglia - astrocyte communication may drive the changes we observe in GFAP expression at the mRNA level in assembloids. Thus, we further assessed DA neurons and astrocytes in our assembloid model using IF. Staining of canonical astrocytic marker GFAP revealed no change in the number of GFAP positive cells, and only a trend towards increased GFAP positive area in assembloids, which did not reach statistical significance (Fig. 4f-g). Staining for tyrosine hydrolase (TH), a dopamine biosynthetic enzyme and canonical DA neuron marker, similarly showed no change in the number of TH positive cells in assembloids. There was however, a significant increase in TH positive area in assembloids, This increased area may reflect increased branching of DA neurons, possibly related to increased maturity of DA circuits within the assembloid (see below). Ultimately, we did not detect changes in the proportion of astrocytes or DA neurons in assembloids compared to hMOs by scRNAseq or IF analysis.

### Integrated microglia impact astrocyte and DA neuron transcriptional signatures in assembloids

We applied pairwise comparisons within the astrocyte and DA neuron clusters in our scRNAseq data to identify DEGs in assembloids compared to hMOs. There were 203 genes upregulated and an additional 153 genes downregulated in the astrocyte cluster derived from assembloids, compared to that derived from hMOs (Fig. 5a). GO analysis of astrocyte DEGs either upregulated or downregulated returned no significantly affected pathways. However, several genes (*KLF6*, *A2M*, *SERPING1*, and *SERPINA3*) whose expression were significantly increased in assembloid astrocytes have been reported to be enriched in reactive mouse astrocytes.^54^ Pairwise comparison revealed 88 DEGs upregulated and an additional 90 DEGs downregulated in DA neurons derived from assembloids versus hMOs (Fig. 5b). GO analysis of upregulated DEGs yielded no significant results, however GO analysis of downregulated DEGs shows that component genes of pathways related to regulation of neural projections and synapse organization are decreased in DA neurons in assembloids (Fig. 5c). These downregulated DEGs include neuronal and synaptic cell adhesion molecules (*CDH4*, *NCAM1*, *NRCAM*, *KIRREL3*, *CADM1*),^55-60^ axon guidance cues (*SEMA6D*, *SLIT1*),^61-63^ and a microtubule motor shown to be important for synaptogenesis (*KIF1A*).^64^ Notably, DA neurons derived from assembloids have significant upregulation of *OTX2*, a transcription factor that plays a key role in DA neurogenesis and marks a subset of adult DA neurons in the ventral tegmental area of the midbrain.^65-69^ Taken together the down-regulation of projection development and guidance terms, and upregulation of *OTX2* may indicate increased maturity of DA neurons in assembloids compared to hMOs. However, the implication of downregulation of synapse organization related terms is less clear. Whether these changes correlate with an actual decrease in synapses, and if they are caused by a defect in neuronal development/maturation, increased synapse removal by microglia, or degenerative loss of synapses, warrants further investigation.

**Figure 5.**
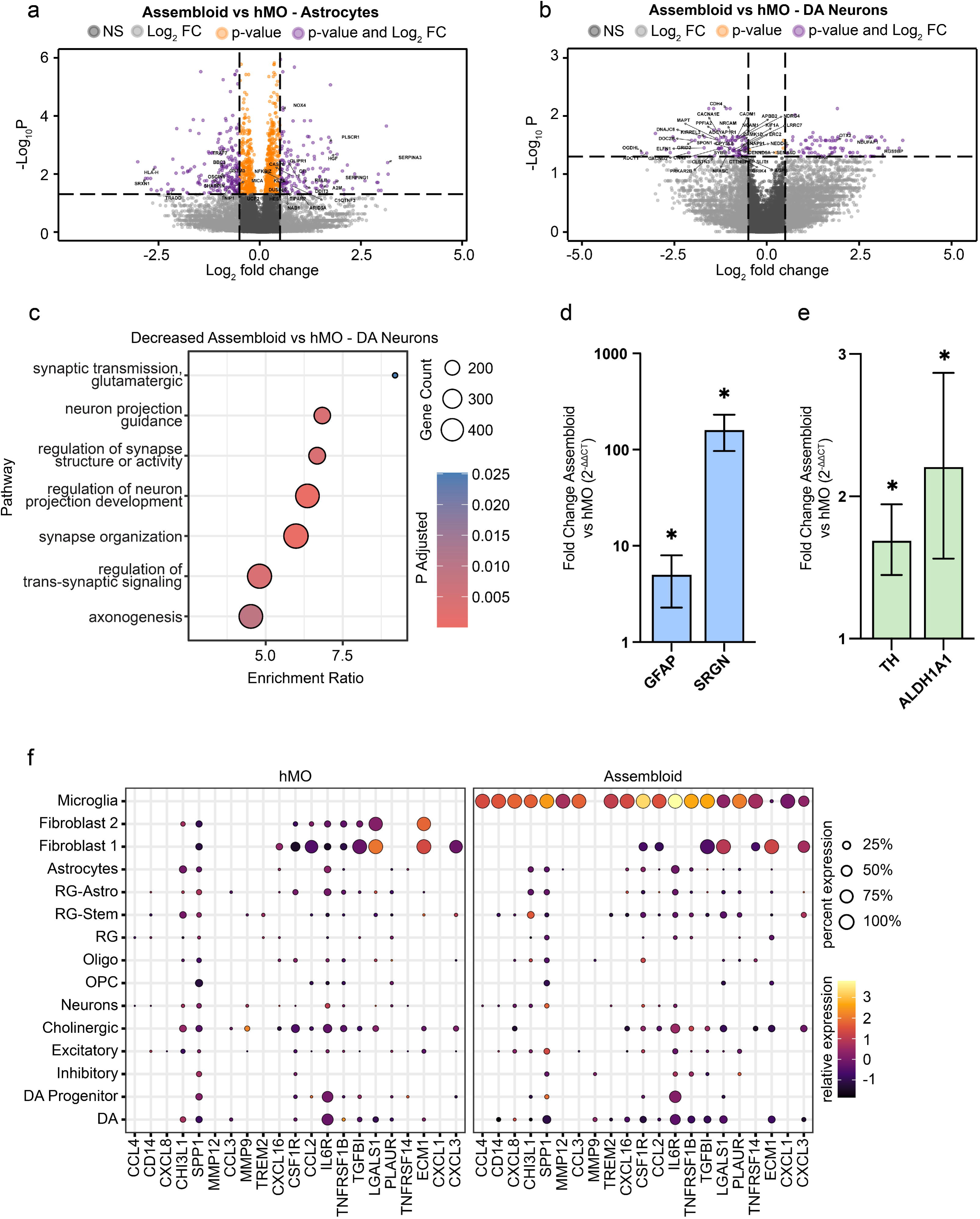
Altered transcriptional signatures of DA neurons and astrocytes in assembloids. **a** DEGs up and downregulated in the astrocyte cluster from assembloids versus hMOs. **b** DEGs up and downregulated in the DA neuron cluster from assembloids versus hMOs. **a-b** Log2FC cut-off = 0.5, p-value cut-off = 0.05 **c** GO terms associated with DEGs decreased in DA neurons derived from assembloids versus hMOs. **d-e** RT-qPCR analysis of **d** reactive astrocyte marker gene GFAP and SRGN, and **e** DA neuron marker gene TH, and ALDH1A1 expression in assembloids normalized to hMOs. n = 4 batches. Unpaired t-test * p ≤ 0.05, ** p ≤ 0.01, *** p ≤ 0.001, **** p ≤ 0.0001 **f** expression of DSPs detected by nELISA in hMO and assembloid scRNAseq data.

To further examine potential changes in astrocyte reactivity and DA neuron maturity, we performed targeted analysis of mRNA expression of relevant genes by RT-qPCR (Fig. 5d). Genes known to be upregulated during astrogliosis, *GFAP* and *SRGN*^45,54,70^ were analyzed. Significant, approximately 10-fold and 200-fold increases were observed in *GFAP* and *SRGN* expression, respectively, in assembloids compared to hMOs. Expression levels of classical DA neuron markers, *TH* and *ALDH1A1* also showed approximately 2-fold and 3-fold increases, respectively, in assembloids compared to hMOs. As we did not observe changes in the number of astrocytes or DA neurons in assembloids compared to hMOs either by scRNAseq or IF, the changes in gene expression detected by RT-qPCR indicate that intMGs drive alterations in the state of these cells. Taken together with DEGs identified in our scRNAseq data, our findings indicate an increased expression of genes related to astrocyte reactivity in assembloids. In DA neurons, we observe increased expression of classical DA genes (*TH*, *ALDH1A1*, and *OTX2*)^67,71^ coupled with decreased projection guidance and synapse organization genes.

We then investigated whether intMG driven alterations in neurons and macroglia might underlie the changes in protein secretion observed by nELISA (Fig. 2b-f). Similar to the cell expression patterns predicted by querying a brain expression data base, our scRNAseq data illustrates that microglia express the highest levels of DSPs increased in assembloids (Fig. 5e). Whereas some DSPs (CHI3L1, LGALS1, PLAUR, ECM1, CXCL1, and CXCL3) were predicted to be expressed in astrocytes and endothelial cells, they are in fact most highly expressed by intMGs in assembloids. Although integration of microglia into assembloids does induce downstream changes in DA neurons and astrocytes, microglia themselves still appear to be the strongest contributor to changes in protein secretion from assembloids.

### Assembloids display altered cell-cell communication networks and an increase in immune-related signaling

Given the role of microglia in orchestrating immune responses, coupled with the changes we have observed in DA neurons and astrocytes in assembloids, we applied the cell-cell communication network analysis program CellChat to our scRNAseq data to further elucidate the impact of intMGs on assembloids.^72,73^ Using a manually curated database of ligand-receptor interactions, CellChat harnesses scRNAseq data to compute the communication probability between two cell clusters based on expression levels of ligands and receptors. Further, CellChat can provide information on a pathway level by calculating the sum of the communication probabilities across all receptor-ligand pairs within a common signaling pathway.^72^ Comparison of the cell-cell communication networks in assembloids versus hMOs reveals four signaling pathways significantly enriched in hMOs, and 52 signaling pathways significantly enriched in assembloids (Fig. 6a). The majority of pathways enriched in assembloids are related to immune function; including regulation of microglial state and function (SPP1/osteopontin, CX3C, GRN, COMPLEMENT),^74-77^ antigen presentation (MHC-I and MHC-II), and cytokine and chemokine signaling (IL-1, IL-2, IL-16, IL-10, TNF, CCL, CXCL). In addition to these immune-related pathways, growth factor signaling pathways are also enriched in assembloids, including VEGF, TGFβ, HGF, GDNF, IGF, EGF, EPGN, and ANGPT. Some of these growth factor ligands have been reported to be released from microglia themselves.^78-82^ To determine which cell types are contributing to the altered signaling network in assembloids, we visualized the relative strength of outgoing signaling (expression of ligands) and incoming signaling (expression of receptors) per pathway across the different cell types in hMOs and assembloids (Fig. 6b-c). intMGs drive novel incoming and outgoing signaling patterns in assembloids compared to hMOs, as evidenced by increased signaling strength in pathways that are absent in hMOs. Notably, these novel signals are not solely sent or received by microglia, but also by the other cells in the assembloid. Approximately half of the pathways significantly enriched in assembloids, as identified in Figure 6a, fit into one of three signaling patterns: i) Expression of both ligand and receptor is restricted to microglia (Fig. 6d); many of these pathways are involved in microglial or peripheral immune cell activation (LIGHT, PD-L1, PD-L2, CD48, CD80, CD86, CD13, IL-10).^83-88^ ii) Ligand expression is restricted to microglia while all other cell types in the organoid express receptors (Fig. 6e). iii) Receptor expression is limited to microglia while ligands are expressed by all cell types of the assembloid - including microglia (Fig. 6f). Of note in this last category is the CX3C pathway. The ligand fractalkine (CX3CL1), mainly expressed by neurons and endothelial cells, binds to the CX3CR1 receptor on microglia to tightly regulate microglial state, and is a commonly used factor in protocols for differentiation of 2D iMGs.^18,77^

**Figure 6.**
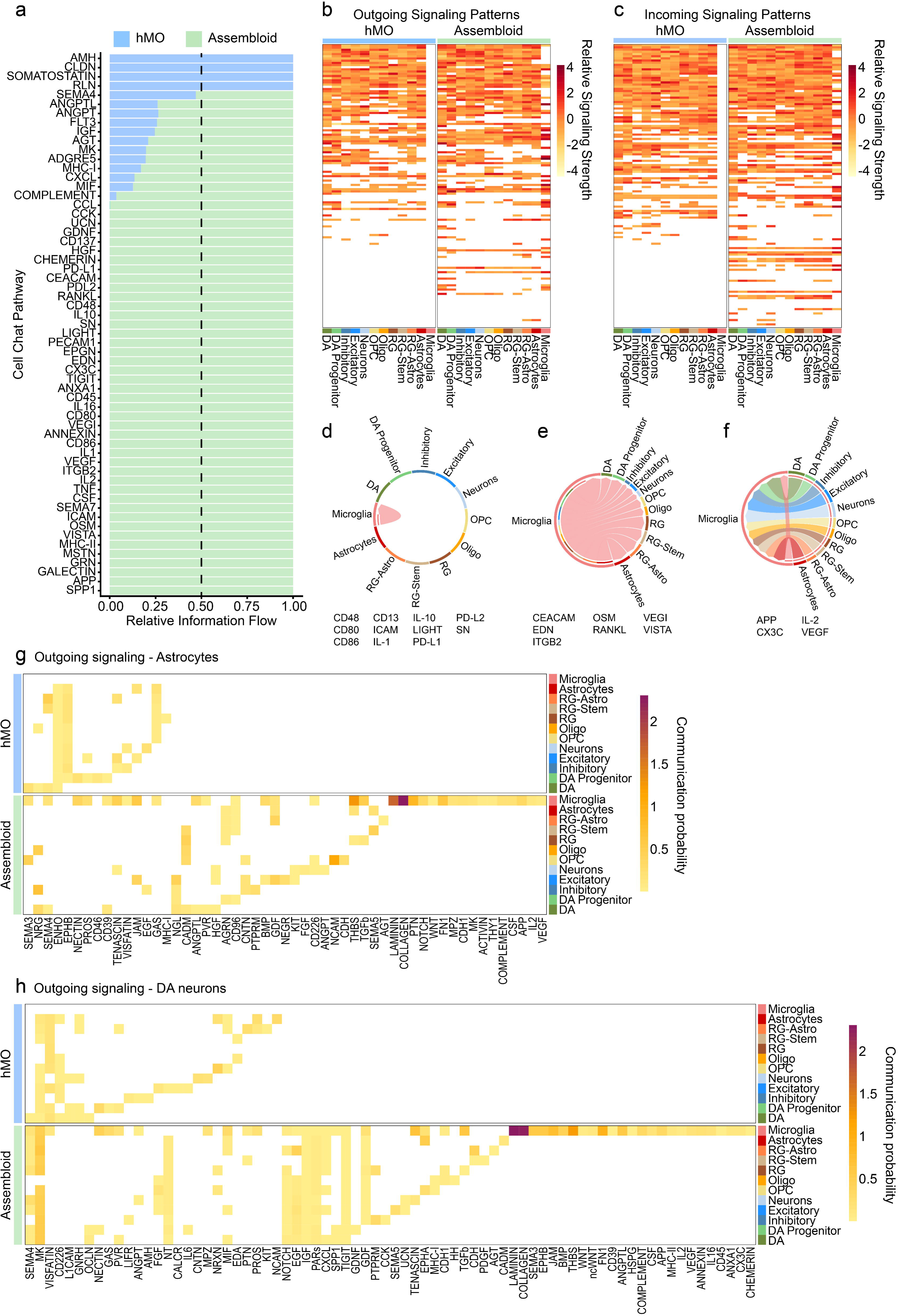
Altered cell-cell communication network in assembloids **a** Relative information flow through CellChat signaling pathways significantly enriched in hMOs and assembloids. Relative signaling strength illustrates altered patterns of **b** outgoing and **c** incoming signaling in assembloids. **d** pattern of microglia to microglia signaling in assembloids, and associated pathways. **e** pattern of microglia to neurons and macroglia signaling in assembloids, and associated pathways. **f** pattern of neuron and macroglia to microglia signaling in assembloids, and associated pathways. **g** Communication probability of outgoing signals from astrocytes in hMOs and assembloids. **h** Communication probability of outgoing signals from DA neurons in hMOs and assembloids.

The remaining pathways enriched in assembloids do not fit into one of these patterns and involve signaling sent from subsets of neurons and macroglia onto microglia and/or other neurons and macroglia. Of particular interest were any changes in astrocyte and DA neuron signaling. We focused on outgoing signals (ligand expression) from astrocytes and DA neurons, plotting pathways whose communication probability was increased or decreased at least two-fold in assembloids compared to hMOs (Fig. 6g-h). The most striking differences between both astrocyte and DA neuron signaling in assembloids compared to hMOs are the pathways whose receptors are expressed by microglia. These pathways are involved in the extracellular matrix (Laminin, collagen, HSPG, Fibronectin/FN1, E-cadherin/CDH1, THY1, and thrombospondin/THBS),^89-91^ development and patterning (semaphorin 3/SEMA3, ephrin b/EPHB, WNT, ncWNT, BMP, VEGF, activin, notch, Midkine/MK, pleiotrophin/PTN, and ANGTL),^61,92-101^ as well as immune and inflammatory processes (COMPLEMENT, CSF, IL-16, CX3C, Chemerin, CD39, MHC-II, IL-2, CD45, ANNEXIN, and ANXA1).^1,76,77,102-108^ Many of these outgoing pathways are received by microglia from both astrocytes and DA neurons. Additionally, both astrocytes and DA neurons send novel outgoing signals to the other cell types in assembloids. In astrocytes the majority of these are related to axon guidance and synapse formation or regulation (NGL, CADM, AGRN, CNTN, PTPRM, GDF, NEGR, NCAM, CDH),^55,56,58-60,109-114^ and are mainly received by the different neuronal cell types and radial glia in the assembloid. Similar changes are seen in DA neurons with novel neurotrophic signaling pathways involved in cell growth and differentiation (NOTCH, EGF, IGF, PARs, GDF),^93,111,115-117^ as well as pathways involved in immune regulation (TIGIT, SPP1, UCN, TENASCIN, MHC-I),^75,118-121^ being received by the various cell types in the assembloid. Overall, CellChat analysis reveals that integration of microglia drives altered cell-cell communication in assembloids. There is a marked increase in immune regulatory and trophic support pathways, further emphasizing the functionality of intMG within assembloids. However, novel signals are also sent from astrocytes and DA neurons onto intMGs and the other cell types present in the assembloid.

## Discussion

Microglia play essential roles in the brain throughout life and during disease. Limitations of animal and 2D cell culture models have contributed to a gap in our understanding of the contributions of microglia to neurodegenerative diseases, including PD. Combination of 3D hMOs with microglia to form assembloids allows for culture of cells in a complex 3D architecture and overcomes the species differences of animal models, providing a promising model for the study of microglia in PD pathogenesis. Here we adapt protocols for hMO and iMG differentiation to generate assembloids.^18,42,43^

We demonstrate that iHPCs integrate into assembloids and mature into intMG that express classical microglia and macrophage markers Iba1, PU.1, and CD68. These intMG display ramified morphology and can persist within assembloids for at least five months. This supports the suitability of our model for future studies in diseases, such as PD, where pathology only arises after long-term organoid culture.^40^

Our model recapitulates the ability of microglia to produce cytokines and chemokines. RT-qPCR and nELISA showed little to no production of cytokine and chemokines by hMOs, even in response to LPS stimulus. LPS signaling is initiated through binding to membrane TLR4 and is known to induce neuroinflammation and gliosis.^122,123^ While TLR4 has been shown to be expressed by both microglia and astrocytes in human brain,^48^ astrocytes lack CD14, a co-receptor necessary for internalization of LPS-bound TLR4, and have been found to have little to no response to LPS in microglia-depleted mice.^45^ We observe a similar lack of response to LPS in hMOs. Conversely, cytokine expression was significantly increased in assembloids following LPS treatment. Additionally, we observe significant increases in a common set of cytokines and chemokines when comparing assembloids to hMOs, regardless of LPS treatment. Many assembloid DSPs are cytokines and chemokines involved in the neuroinflammatory response, however, secretion of a number of proteins related to ECM maintenance was also increased. This likely reflects the adaptation of microglia to the complex 3D environment of the assembloid. The largely overlapping signature of protein release in assembloids, regardless of LPS treatment, illustrates that the integration of microglia alone is enough to induce inflammatory signaling in assembloids. This may reflect a base-line level of microglial activity in response to DAMPs and signaling from other cell types present within the assembloid, that can then be further potentiated with LPS. However, the greatest differences in protein release are between the different culture types assessed – 2D iMGs, hMOs, and assembloids. Similar results, including increased secretion of chemokines CCL2, CCL3, and CCL4 from microglia-hMO assembloids, have been reported.^41^ LPS treatment of assembloids also resulted in an increase in GFAP expression that was absent in hMOs, underscoring the functionality of intMG and their ability to recapitulate the inflammatory signaling from microglia to astrocytes observed *in vivo*.^45^

To more deeply assess the impact of the complex mix of cell types within the 3D environment of an assembloid on intMG, and vice – versa, we performed scRNAseq. Although, intMG expressed the same microglia markers as 2D microglia, they displayed a unique transcriptional signature compared to both 2D microglia and the neurons and macroglia derived from hMOs and assembloids. Differentiation in the 3D environment of the assembloid led to downregulation of a large number of genes involved in RNA and ribosome processing, cell cycle, and proliferation pathways characteristic of immature microglial populations found in embryonic and early post-natal mouse brains.^49^ These changes are coupled with upregulation of genes involved in pathways regulating cell activation and adhesion, congruent with a transcriptional program enriched in cytokine responsive microglia and an immune-alerted population enriched in mouse midbrain.^50,51^ Similar alterations to microglial transcriptional state, including upregulation of a cytokine-associated signature, have been reported upon integration in cortical brain organoids.^31,36^ Microglial transcriptional state is known to be highly dependent on environment; here we demonstrate for the first time that microglia differentiated within a microglia-hMO assembloid adopt a less proliferative, more inflammatory state compared to those differentiated in 2D.

We observed no changes in the proportion of neurons and macroglia within assembloids. However, several of our findings indicate changes in the state of astrocytes and DA neurons in assembloids compared to hMOs. Several genes known to be enriched in reactive astrocytes were found to be upregulated in assembloid-derived astrocytes. DA neurons derived from assembloids had significantly decreased expression of genes involved in regulation of neural projection formation and synapse organization, and increased expression of DA marker genes (*TH*, *ALDH1A1*, and *OTX2*). Integration of microglia into hMOs, by Sabate-Soler and colleagues, led to downregulation of genes involved in synapse remodeling and axon guidance, and electrophysiological changes indicative of more mature neuronal circuitry.^41^ Although the specific genes queried by Sabate-Soler and colleagues were not identified as significantly downregulated in DA neurons derived from assembloids in our scRNAseq data, we did observe significant downregulation of genes functioning in similar synapse organization and axon guidance pathways. Study of the electrophysiological properties of our assembloid model, including use of pharmacological agents targeting DA circuits, is warranted and will provide better understanding of how the changes we observe may influence neuronal maturation and circuitry.

Analysis of our scRNAseq data illustrates that integration of microglia into assembloids leads to significantly increased expression of ligand-receptor pairs involved in inflammatory and growth factor signaling pathways. This demonstrates that two main microglial functions: induction and regulation of inflammatory signaling, and trophic support, are occurring in assembloids. Further, integration of microglia into assembloids leads to altered outgoing signaling from astrocytes and DA neurons, including patterning and immune-regulatory signals received by microglia, and neurotrophic and axon guidance signaling received by the various neurons and macroglia of the assembloid. The increased maturity and immune-responsive state of intMGs compared to 2D microglia is likely mediated in part by the many signaling pathways received from the other cell types in the assembloid. Together these signaling network changes reflect the broad impact of intMG on assembloids, and vice-versa.

Assembloids provide a promising model for investigation of the contribution of microglia and neuroinflammation to PD. intMG are functional, with a more mature and inflammation-responsive transcriptional state than 2D microglia and induce increased trophic and inflammatory signaling in assembloids as a whole. The assembloid model described and characterized here should be further harnessed to study PD pathogenesis mechanisms reported to impact or be impacted by immune function. Generation of assembloids from PD patient-derived iPSC lines carrying mutations that induce α-synuclein pathology, mitochondrial and autophagic-lysosomal pathway dysfunction would be of particular interest. Our assembloid model has the potential to help delineate the impact of microglial, astrocytic, and cell-autonomous mechanisms on DA neuron death in PD.

## Methods

### iPSC cell culture

The use of human iPSCs and iPSC-derived cells in this research was approved by the McGill University Research Ethics Board (IRB Study Number A03-M19-22A). A healthy control iPSC line was used to generate all cultures used in this study. This line (IPSC0063) is registered with the hPSCReg repository (https://hpscreg.eu/cell-line/CBIGi001-A). This line underwent whole genome sequencing to preclude the presence of any disease associated variants, and was subjected to quality control measures including karyotyping, assessment of pluripotency marker expression, and regular mycoplasma testing. iPSCs were cultured on matrigel coated dishes in mTESR1. iPSCs were passaged a minimum of two times prior to differentiation of hMOs, iMGs, or assembloids. All iPSC culture and differentiation reagents are listed in supplementary table 1.

### hMO generation

hMOs were generated following a previously published protocol.^42,43^ Briefly, 10 000 iPSCs in single cell suspension were seeded per well in a 360-well ULA-coated EB DISK (eNUVIO) in neuronal induction media (1:1 DMEM+Anti:Neurobasal, 1:100 N2, 1:50 B27 without vitamin A, 1% Glutamax, 1% MEM-NEAA, 5 µM 2-mercaptoethanol, 1 µg/mL Heparin, 10 µM SB431542, 200ng/mL Noggin, 0.8 µM CHIR99021, 10 µM Y-27632). iPSC - containing EB disks were centrifuged in 6 well plates at 1200 rpm for 10 minutes prior to incubation at 37°C and 5% CO2. Embryoid bodies (EBs) form after two days of culture, and Y-27632 is withdrawn from the media. After an additional two days, a full media change to midbrain patterning media (1:1 DMEM+Anti:Neurobasal, 1:100 N2, 1:50 B27 without vitamin A, 1% Glutamax, 1% MEM-NEAA, 5 µM 2-mercaptoethanol, 1 µg/mL Heparin, 10 µM SB431542, 200 ng/mL Noggin, 0.8 µM CHIR99021, 100 ng/mL SHH, 100 ng/ml FGF-8) is performed. Three days later, EBs are embedded in a reduced growth factor Matrigel scaffold and a full media change to tissue growth media (Neurobasal, 1:100 N2, 1:50 B27 without vitamin A, 1% Glutamax, 1% MEM-NEAA, 5 µM 2-mercaptoethanol, 200 ng/mL laminin, 2.5 µg/mL insulin, 100 ng/mL SHH, 100 ng/ml FGF-8, 1X Penicillin-Streptomycin) was performed. The next day, EBs were transferred to ultra-low attachment 6-well plates containing final differentiation media (Neurobasal, 1:100 N2, 1:50 B27 without vitamin A, 1% Glutamax, 1% MEM-NEAA, 5 µM 2-mercaptoethanol, 10 ng/mL BDNF, 10 ng/mL GDNF, 100 µM ascorbic acid, 125 µM db-cAMP, 1X Penicillin-Streptomycin), and maintained on orbital shakers at 70 rpm at 37°C and 5% CO2. hMOs were cultured for two months with 50% media changes performed three times a week before integration of microglia. All iPSC culture and differentiation reagents are listed in supplementary table 1.

### 2D iMG differentiation

iMGs were differentiated following a previously published protocol.^18^ Briefly, iPSCs were seeded at multiple densities onto Matrigel coated 6-well plates in mTESR1. iHPC differentiation was performed using the STEMCELL Technologies STEMdiff Hematopoietic Kit. iHPCs were harvested on days 10 and 12 of differentiation and re-plated onto Matrigel coated 6-well plates in microglial differentiation media (DMEM-F12, 1:50 Insulin-transferrin-selenite, 1:25 B27, 1:200 N2, 1% Glutamax, 1% MEM Non-essential amino acids, 400 μM monothioglycerol, and 5 μg/mL human insulin) + three cytokine cocktail (25 ng/mL M-CSF, 100 ng/mL IL-34, and 50 ng/mL TGF-β). Cells were supplemented with microglia differentiation media + three cytokine cocktail every other day. A full media change was performed on day 12 post iHPC re-plating with microglia differentiation media + three cytokine cocktail. Another full media change was performed on day 25 post iHPC re-plating with microglia differentiation media + five cytokine cocktail (25 ng/mL M-CSF, 100 ng/mL IL-34, 50 ng/mL TGF-β, 100 ng/mL CX3CL1, and 100 ng/mL CD200). Cells were supplemented with microglia differentiation media + five cytokine cocktail on days 27 and 29, and used for experiments on day 30. Three and five cytokine cocktails were always added fresh immediately before media supplementation or change. All iPSC culture and differentiation reagents are listed in supplementary table 1.

### Generation of intMG-hMO assembloids

To generate assembloids iHPCs were generated using the STEMCELL Technologies STEMdiff Hematopoietic Kit, and harvested on days 10 and 12 of differentiation as described above. Instead of being re-plated for iMG differentiation iHPCs were added into ultra-low attachment 6-well plates with 2 month old hMOs in assembloid media (DMEM-F12, 1:50 Insulin-transferrin-selenite, 1:25 B27, 1:200 N2, 1% Glutamax, 1% MEM Non-essential amino acids, 400 μM monothioglycerol, 5 μg/mL human insulin, 25 ng/mL M-CSF, 100 ng/mL IL-34, 10 ng/mL BDNF, 10 ng/mL GDNF, 100 µM ascorbic acid, and 1X Penicillin-Streptomycin) at a ratio of 50 000 iHPCs per hMO. M-CSF and IL-34 were always added fresh to assembloid media immediately before use. Cultures were maintained on orbital shakers at 70 rpm at 37°C and 5% CO2. To allow iHPC colonization of the hMO tissue, media was supplemented with assembloid media two days after iHPC addition, a 50% media change was performed four days after iHPC addition, and then 50% media changes with assembloid media were performed three times per week. Assembloids were harvested and used for downstream experiments 30 days after the addition of iHPCs. All iPSC culture and differentiation reagents are listed in supplementary table 1.

### Cryosectioning and immunofluorescence staining of hMOs and assembloids

hMOs and assembloids were fixed in 4% paraformaldehyde (ThermoFisher 28908) in PBS overnight at 4°C. Fixed hMOs and assembloids were then equilibrated in 20% sucrose in PBS for 2-3 days, until no longer floating. hMOs and assembloids were then embedded in optimal cutting temperature compound (ThermoFisher 23730571) and stored at - 80°C. Embedded hMOs and assembloids were sectioned at 10 μm thickness using a cryostat, and mounted onto Superfrost positively charged glass slides (ThermoFisher 12-550-15). Slides were stored at -20°C. For immunofluorescence staining, slides were rehydrated in PBS for 15 minutes in a slide staining jar on an orbital shaker at room temperature. Slides were then incubated in permeabilization/blocking solution (PBS containing 5% normal donkey serum (Millipore S30), 0.05% bovine serum albumin (Wisent 800-095-EG), and 0.2% Triton-X 100 (ThermoFisher BP151-500)) for one hour at room temperature. Slides were then incubated with primary antibodies diluted in permeabilization/blocking solution overnight in a humidified chamber at 4°C. Slides were washed 3 x 5 minutes in PBS in a slide staining jar on an orbital shaker at room temperature, followed by incubation with fluorescently coupled secondary antibodies diluted in permeabilization/blocking solution in a humidified chamber at room temperature for one hour. Slides were washed as above, then incubated with a 1:5 000 dilution of Hoechst 33342 in PBS for 10 minutes at room temperature. After 2 x 5 minute final washes in PBS, coverslips were mounted using aqua-poly/mount (PolySciences 18606-20). Slides were imaged using an oil-immersion 40X objective on a Leica TCS SP8 confocal microscope. Three sections were imaged per organoid. CellProfiler was used to quantify staining area and cell number, Hoechst 33342 area and counts were used for normalization. Antibodies are listed in supplementary table 2.

### CUBIC tissue clearing and 3D imaging

Assembloids were cleared using the CUBIC (Clear, Unobstructed Brain/Body Imaging Cocktails and computational analysis) method.^124^ Briefly, fixed assembloids were immersed in CUBIC R1 tissue clearing solution (25% urea (Sigma-Aldrich STBH2864), 25% Quadrol (N, N, Ń, Ń-tetrakis (2-hydroxy-propyl) ethylenediamine) (Sigma-Aldrich 102-60-3) and 15% Triton X-100 (Bioshop 4C33340) in distilled water) at 37℃ until full transparency, washed in PBS and incubated in Permeabilization/Blocking solution (5% Glycine 2M (Bioshop GLN001.5), 5% Normal Donkey Serum (Millipore S30-100mL), 0.6% Bovine Serum Albumin (80mg/mL MultiCell 800-095-CG)), 0.2% Triton X-100 (Millipore TX1568-1) in PBS overnight at 4℃. The samples were stained with primary antibodies for two days at 37℃, washed, counterstained with Hoechst 33342 and washed before incubation in 20% sucrose solution (Fisher bioreagents BP220-1) until samples precipitated. Finally, the assembloids were immersed overnight at 37℃ in CUBIC R2 tissue clearing solution (1% triethanolamine (FisherScientific 22-010-1223), 50% sucrose (Fisher bioreagents 57-50-1) and 25% of urea (Sigma-Aldrich STBH2864) in distilled water), before imaging. Each step was performed under gentle shaking (∼100 rpm). Confocal images of cleared assembloids (2 channels, 2 fields/organoid, 100 stacks/field at 2µm increment) were acquired with PerkinElmer Opera Phenix equipped with a 20x water objective NA1.0.

### RNA extraction and RT-qPCR

2D iMGs, hMOs, and assembloids were treated with 100 ng/mL LPS (ThermoFisher 00-4976-93) for 24 hours. Samples were lysed in trizol (ThermoFisher 15596026) and RNA extraction was performed using the Qiagen RNeasy kit (74106). RNA concentrations were determined using a nanodrop spectrophotometer, and a reverse transcription reaction was set up using 500 ng of RNA, M-MLV reverse transcriptase (ThermoFisher 28025013), random hexamers (ThermoFisher N8080127), RNase inhibitor (ThermoFisher N80801), and dNTPs (New England Biolabs N0447L) diluted in the 5X First strand buffer + DTT supplied with the M-MLV reverse transcriptase. Reverse transcription reactions were then incubated in a Bio-Rad T100 thermal cycler as follows: 42°C for 60 minutes, 75°C for 10 minutes. Resulting cDNA was then stored short term at -20°C before use in RT-qPCR. RT-qPCR was performed using taqman assays (ThermoFisher) on a quantstudio 3 real-time PCR system (ThermoFisher). Data were analyzed using the comparative threshold method, with normalization to GAPDH. Taqman assays are listed in supplementary table 3.

### nELISA secreted protein quantification

Conditioned media from LPS-treated and non-treated 2D iMGs, hMOs, and assembloids used for RT-qPCR described above was collected, spiked with bovine serum albumin (Wisent 800-095-EG) to a final concentration of 0.1% and stored at -80°C. nELISA and determination of protein concentration in pg/mL was performed by Nomic (5333 Casgrain Ave, Suite 401, Montréal, Québec, Canada, H2T 1X3) utilising conditioned media from two distinct batches of iMGs, and four of hMOs/assembloids.^47^ In cases where the fluorescence intensity was out of the range, concentrations corresponding to upper and lower limits of detection were used as appropriate. For pairwise comparison, concentration fold change was calculated, and significance was determined by student’s t-test of fluorescence intensity values. Secreted proteins with a fold change greater than or equal to 2 and a p-value less than or equal to 0.05 were considered significant.

### Single cell RNA sequencing – sample preparation and sequencing

Ten hMOs and ten assembloids were each pooled and dissociated using TrypLE express without phenol red (Gibco 12604-013) in a GentleMACS M-Tube (Miltenyi Bitoec 130-093-236) on the automated GentleMACS Octo Heated dissociation device (Miltenyi Biotec) set to spin for 24 minutes at 20 rpm, followed by one minute at 197 rpm, both at 37°C. After dissociation TrypLE was quenched by addition of PBS spiked with RNase inhibitor (ThermoFisher N80801) and DNase (Qiagen 79254). Cell suspensions were transferred using low bind pipette tips to a 15 mL falcon tube through a 30 μm mesh filter (Miltenyi Biotec 130-041-407). 2D iMGs from three wells of a 6 well plate were detached by scraping in PBS and pooled. iMGs, and hMO and assembloid cell suspensions were washed three times in PBS spiked with RNase inhibitor and DNase, with centrifugation at 350 x g for 5 minutes at 4°C to pellet cells between washes. Cells were then fixed using the Parse Biosciences Evercode Cell Fixation v3 Kit, following the manufacturer’s protocol. Fixed cells were stored at -80°C. Fixed cells were thawed, counted, and processed using the Parse Biosciences Evercode WT v3 Kit. The resulting libraries underwent quality control and sequencing at the McGill Genome Centre (740 Dr. Penfield Avenue, Montreal, Quebec H3A 0G1 Canada) on an Illumina NovaSeqXPlus at a depth of 50 000 reads per cell.

### Single cell RNA sequencing – data analysis Trailmaker and Seurat

Fastq files were then processed on Parse Biosciences Trailmaker pipeline (v1.5.1), the resulting filtered count matrices were analyzed using the Seurat (v5.4) computational workflow. Briefly, cells with high mitochondrial content (>20% of reads), low UMI counts (<100), low distinct gene counts (<100), and putative doublets were removed. The Seurat object containing all 2D iMG, hMO, and assembloid data was subsequently normalized and scaled, and principal component analysis (PCA) was preformed using the top 2000 highly variable features. The top 10 principal components were used to preform Louvain clustering and Uniform Manifold Approximation and Projection for Dimension Reduction (UMAP) for visualization of the data. Here, clusters were defined based on sample identity (iMG, hMO, or hMO + intMG) and expression of microglial markers. Similarity to transcriptional programs defined by Fumagalli and microglial clusters identified by Uriarte Huarte was assessed using the AddModuleScore function in seurat. The hMO and hMO + intMG samples were then isolated and underwent anchor-based integration to account for variations due to different cell populations (i.e. the presence of microglia in hMOs + intMG relative to hMOs). The resulting data was re-clustered using the above workflow and we preformed Adaptively-thresholded Low Rank Approximation (ALRA) to impute dropouts and account for false zero expressions. A clustering resolution of 1 was chosen and clusters were annotated based on their expression of characteristic cell type markers. DA neurons had high expression of *TH*, *LMX1A*, *EN1*, *CALB1*, and *OTX2*, in addition to *FOXA1*, *FOXA2*, *NDNF*, *ANXA1*, and *DDC* – which were also expressed in the DA progenitor cluster. Inhibitory, excitatory, cholinergic, and broad neuron clusters were marked by expression of *STMN2* and *SYP.* Inhibitory neurons were differentiated by expression of *GAD1*, *GAD2*, *SLC32A1*, and *SOX14*, excitatory neurons by expression of *SLC17A6*, *SLC17A7*, and *NEUROD2*, and cholinergic neurons by expression of *ACHE*, and *CHAT*. Oligodendrocytes (Oligo) and Oligodendrocyte precursors (OPC) were marked by high expression of *OLIG1*, *OLIG2*, and *PLP1*; with additional expression of proliferation markers *PCNA*, and *MKI67* differentiating OPC. Radial glia (RG) cells, in addition to two related clusters of stem cell – like RG (RG-Stem) and RG – astrocyte transition cells (RG-Astro), expressed *PAX6*, *GLI3*, and *HES5*. RG-stem also expressed proliferation markers *PCNA* and *MKI67*. The RG-Astro and astrocyte clusters expressed *GFAP*, *ALDH1L1*, and *SOX9*. Fibroblasts were identified by high expression of *COL15A1*, *COL3A1*, *COL1A3*, *MGP*, and *MMP12*. Finally, the microglia cluster was annotated based on high expression of *AIF1* (IBA1), *SPI1* (PU.1), *P2RY12*, and *MERTK*. Permutation tests were performed using the scProportionTest package in R. Differentially expressed genes (DEGs) were extracted using Seurat’s FindMarkers function (MAST test) With an FDR threshold of 0.05 and Log2FC threshold of 1 for comparison of 2D vs 3D microglia, and 0.5 for comparison within astrocyte and DA neuron clusters derived from hMOs and assembloids.

### CellChat

The cell-cell communication network of hMOs and assembloids was then assessed using the CellChat (v1.6.1) computational workflow for comparative analysis of samples with different cell populations. Briefly, hMO and assembloid data was first processed separately. Cell clusters with fewer than 10 cells were excluded. Using the CellChat human database trimean ligand-receptor communication probabilities were calculated and summed to generate pathway level communication probabilities. Pathway level communication probabilities were then further summarized to generate the aggregated cell-cell communication network. Network centrality scores were then calculated to determine major signaling targets and sources. The CellChat objects corresponding to the hMO and assembloid samples were then lifted and merged for comparative analysis. The rankNet function was used to determine pathways significantly enriched in the assembloid versus hMO by comparing the strength of pathway information flow. CellChat visualization functions were then used to generate heatmaps and chord diagrams representing communication probability and centrality scores of these pathways.

### Gene Ontology Over representation analysis

DSPs detected by nELISA as well as DEGs detected by scRNAseq were analyzed for Gene Ontology terms related to biological processes and molecular function using the WebGestalt toolkit.

### Query of brain RNA sequencing data set

Human brain RNA sequencing data used to predicted cell type expression of DSPs was downloaded from brainrnaseq.org.

### Statistical Analysis

All statistical analysis was performed using GraphPad Prism version 10.4.1 for MacOS, (GraphPad Software, Boston, Massachusetts USA, www.graphpad.com). All analysis includes data from 2-4 distinct differentiations/batches of iMGs, hMOs and assembloids. Throughout this work n = number of batches. For immunofluorescence analysis, n = 3 batches with 2-3 hMOs or assembloids analyzed per batch, and three sections analyzed per organoid. The mean of sections per hMO/assembloid was calculated, and outliers were removed using the ROUT method for outlier detection (Q = 1%). P-values were calculated using an unpaired, two-tailed t-test, counting each organoid as a replicate. For RT-qPCR analysis of cytokine and GFAP expression n = 2 batches of iMGs, and 4 batches of hMOs/assembloids with 2-3 wells of iMGs and 3-5 hMOs/assembloids pooled per batch for RNA extraction. RT-qPCR reactions were performed in duplicate and the CT mean of these technical replicates was then normalized to GAPDH, and further normalized to untreated controls (2^-ΔΔCT^). P-values were calculated using a two way ANOVA, with Šídák’s multiple comparisons test, counting each batch as a replicate. For RT-qPCR analysis of DA neuron and astrocyte markers n = 4 batches of hMOs/assembloids with 3-5 hMOs/assembloids pooled per batch for RNA extraction. RT-qPCR reactions were performed in duplicate and the CT mean of these technical replicates was then normalized to GAPDH, assembloid samples were then further normalized to hMOs (2^-ΔΔCT^). P-values were calculated using an unpaired, two-tailed t-test, counting each batch as a replicate. All data is presented as mean ± standard error mean. ns p > 0.05, * p ≤ 0.05, ** p ≤ 0.01, *** p ≤ 0.001, **** p ≤ 0.0001

## Author Contributions

EJM and GD contributed equally to this work. EJM generated iHPCs and microglia, generated assembloids, performed immunofluorescence experiments, analysed immunofluorescence data, nELISA data, and scRNAseq data, designed experiments, and prepared the manuscript. GD generated hMOs and assembloids, performed immunofluorescence experiments, qPCR experiments, analysed qPCR and scRNAseq data, designed experiments, and prepared the manuscript. AO, JL, and NF performed immunofluorescence and qPCR experiments, and analysed qPCR data. JS aided in dissociation of organoids for scRNAseq. TMG processed samples for scRNAseq and aided in scRNAseq data analysis. MBT, PL, and MHB performed tissue clearing and 3D imaging. TMD designed experiments. EAF supervised the project, designed experiments, and prepared the manuscript.

## Acknowledgements

Thanks to Vanessa Omana for support with sample processing for scRNAseq, and cryostat training and management. Thanks to Rhalena Thomas for training and advice on scRNAseq and nELISA data analysis. Thanks to Marie-France Dorion for training and advice on differentiation of iPSC-derived microglia. Thanks to Nguyen-Vi Mohamed and Meghna Mathur for training and advice on generation of hMOs. Thanks to Xiuqing Chen for quality control and facilitating access to iPSCs. Thanks to the Neuro Microscopy Core Facility for confocal microscope management, maintenance, and training, and to Camille Lacarrière for CellProfiler training.

Thanks to Wolfgang Reintsch for high content microscope management, maintenance, and training. Thanks to Nomic (5333 Casgrain Ave, Suite 401, Montréal, Québec, Canada, H2T 1X3) for performing nELISA and support with data analysis. Thanks to Frédérique Larroquette for facilitation of nELISA.

EJM has been supported by a Fonds de Recherche du Québec-Santé Doctoral Fellowship and Jeanne Timmins Costello Fellowship awarded by the Integrated program in Neuroscience at McGill University. GD has been supported by a McGill Healthy Brains for Healthy Lives (HBHL) graduate student fellowship, McGill Jeanne Timmins Costello Studentship, and the Parkinson Canada Graduate Student Award. EAF is supported by a Canada Research Chair (Tier 1) in Parkinson’s disease. This work was supported by a project grant from the Canadian Institutes of Health Research (PJT-195804) and a Fonds d’Accéleration des Collaborations en Santé grant (Fon-FACS-013) from CQDM/MEI.

## Data Availability

scRNAseq data generated during this study has been deposited on zenodo (<u>10.5281/zenodo.19410802</u>). Full cluster marker and DEG lists from scRNAseq analysis, along with nELISA fluorescence intensity and protein concentration data are included in supplementary file 1.

## Figure Captions

**Supplementary Figure 1.**
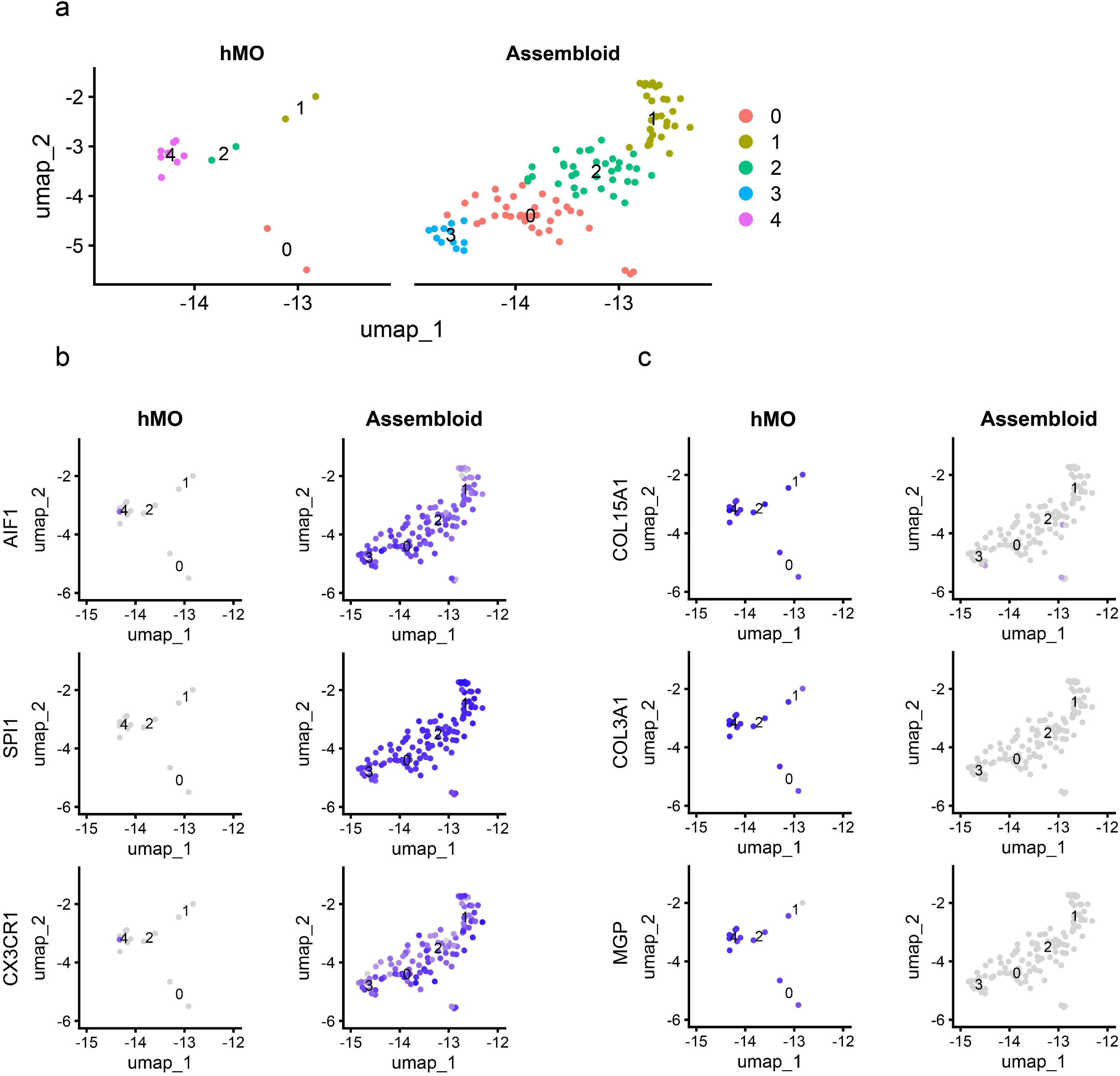
Re-annotation of hMO cells in microglia cluster. **a** UMAP representation showing cells present in microglia subclusters in hMO and assembloids. **b** Expression of microglial marker genes across microglial subclusters in hMOs and assembloids. **c** Expression of fibroblast marker genes across microglial subclusters in hMOs and assembloids.

**Supplementary movies 1-5** 3D imaging reveals ramified morphology of integrated microglia. Representative fields of 3D reconstruction of IBA1 immunofluorescence staining of cleared assembloids in green and counterstaining with Hoechst.

