## Supplemental Tables for "Integration of iPSC-derived microglia into human midbrain organoids enhances microglial maturation and inflammatory signaling"

**Supplementary Table 1 – iPSC culture and differentiation reagents**

| **Reagent** | **Supplier** | **Catalog number** |
| --- | --- | --- |
| 2-mercaptoethanol | Merck | 8057400005 |
| Antibiotic-Antimycotic | Gibco | 15240-062 |
| Ascorbic acid | Sigma Aldrich | A5960 |
| B27 | Gibco | 17504044 |
| B27 without vitamin A | Gibco | 12587010 |
| BDNF | Peprotech | 450-02 |
| Bovine Serum Albumin | MultiCell | 800-095-CG |
| CD-200 | Bon Opus Biosciences | C311 |
| CHIR99021 | Selleckchem | S2924 |
| CX3CL1 | Peprotech | 300-31 |
| db-cAMP | Carbosynth | ND07996 |
| DMEM F12 | Gibco | 10565-018 |
| DMEM F12 HEPES | Wisent | 319-085-CL |
| EB Disc 360 well ULA-Coated | eNUVIO | eN-eb360u-001 |
| FGF-8 | Peprotech | 100-25 |
| GDNF | Peprotech | 450-10 |
| Glutamax | Gibco | 35050061 |
| Heparin | Sigma Aldrich | H3149 |
| Human Insulin | Sigma | I2643 |
| IL-34 | Peprotech | 200-34 |
| Insulin-transferrin-selenite | Gibco | 41400045 |
| Laminin | Sigma Aldrich | L2020 |
| M-CSF | Peprotech | 300-25 |
| Matrigel | Corning | 8774552 |
| Matrigel reduced growth factor | Corning | 356230 |
| MEM Non-essential Amino Acids | Wisent | 321-011-EL |
| Monothioglycerol | Sigma | M6145 |
| mTESR1 | StemCell Technologies | 85850 |
| N2 | Gibco | 17504048 |
| Neurobasal | Life Technologies | 21103-049 |
| Noggin | Peprotech | 120-10C |
| Penicillin-Streptomycin | Wisent | 450-200-EL |
| SB431542 | Selleckchem | S1067 |
| SHH | Peprotech | 100-45 |
| STEMdiff™ Hematopoietic Kit | StemCell Technologies | 5310 |
| TGFβ1 | Peprotech | 100-21 |
| Ultra low attachment 6-well plates | Corning | 3471 |
| Y-27632 | Selleckchem | S1049 |

**Supplementary Table 2 – Antibody information**

| **Antibody** | **Supplier** | **Catalog number** | **Dilution** | **IF Application** |
| --- | --- | --- | --- | --- |
| Anti IBA1 rabbit polyclonal | Wako | 019-19741 | 1:1000 | sectioned tissue |
| Anti IBA1 chicken recombinant monoclonal | Synaptic systems | 234 009 | 1:1000 | sectioned tissue |
| Anti PU.1 rabbit polyclonal | Cell signaling technology | 2266 | 1:500 | sectioned tissue |
| Anti CD68 mouse recombinant monoclonal | Abcam | ab955 | 1:1000 | sectioned tissue |
| Anti GFAP rabbit polyclonal | Agilent dako | Z033429-2 | 1:1000 | sectioned tissue |
| Anti TH chicken polyclonal | Millipore Sigma | AB9702 | 1:500 | sectioned tissue |
| AlexaFluor 555 Anti IBA1 rabbit monoclonal | Cell signaling technology | 36618S | 1:500 | cleared tissue and 3D imaging |
| AlexaFluor 647 donkey anti rabbit | Invitrogen | A32795 | 1:500 | sectioned tissue |
| AlexaFluor 488 donkey anti chicken | Invitrogen | A78948 | 1:500 | sectioned tissue |
| AlexaFluor 555 donkey anti chicken | Invitrogen | A78949 | 1:500 | sectioned tissue |
| AlexaFluor 488 donkey anti mouse | Invitrogen | A21202 | 1:500 | sectioned tissue |

**Supplementary Table 3 – Taqman assays**

| **Target** | **Supplier** | **Catalog number** |
| --- | --- | --- |
| IL1β | ThermoFisher | Hs01555410_m1 |
| IL6 | ThermoFisher | Hs00174131_m1 |
| TNFα | ThermoFisher | Hs00174128_m1 |
| GFAP | ThermoFisher | Hs00909233_m1 |
| AMIGO2 | ThermoFisher | Hs00827141_g1 |
| SRGN | ThermoFisher | Hs01004159_m1 |
| S100A10 | ThermoFisher | Hs00237010_m1 |
| TH | ThermoFisher | Hs00165941_m1 |
| ALDH1A1 | ThermoFisher | Hs00946916_m1 |
| SLC6A3 | ThermoFisher | Hs00997374_m1 |
| GAPDH | ThermoFisher | Hs02786624_g1 |
